# Spatially resolved transcriptome profiles of mammalian kidneys illustrate the molecular complexity of functional nephron segments, cell-to-cell interactions and genetic variants

**DOI:** 10.1101/2020.09.29.317917

**Authors:** Arti M. Raghubar, Duy T. Pham, Xiao Tan, Laura F. Grice, Joanna Crawford, Pui Yeng Lam, Stacey B. Andersen, Sohye Yoon, Monica S.Y. Ng, Siok Min Teoh, Samuel E. Holland, Anne Stewart, Leo Francis, Alexander N. Combes, Andrew J. Kassianos, Helen Healy, Quan Nguyen, Andrew J. Mallett

## Abstract

Understanding the molecular mechanisms underlying mammalian kidney function requires transcriptome profiling of the interplay between cells comprising nephron segments. Traditional transcriptomics requires cell dissociation, resulting in loss of the spatial context of gene expression within native tissue. To address this problem, we performed spatial transcriptomics (ST) to retain the spatial context of the transcriptome in human and mouse kidneys. The generated ST data allowed spatially resolved differential gene expression analysis, spatial identification of functional nephron segments, cell-to-cell interaction analysis, and chronic kidney disease-associated genetic variant calling. Novel ST thus provides an opportunity to enhance kidney diagnostics and knowledge, by retaining the spatial context of gene expression within intact tissue.

## Background

The mammalian kidney is composed of functional nephron segments, including glomeruli, tubules, collecting ducts and microvasculature, spanning the cortical and medullary regions [1,2]. The nephrons maintain homeostasis of body fluids, electrolyte and acid-base balance, and the excretion of metabolic waste products [1,3–5]. The spatial organisation of nephrons facilitates the homeostatic function of the mammalian kidney. However, to date transcriptome studies of nephrons have utilised single-cell and/or single-nucleus RNA-sequencing (scRNA-seq/snRNA-seq), which require manipulation of tissue, including cell dissociation, resulting in the loss of crucial spatial information [6–13].

Unlike scRNA-seq and snRNA-seq, ST-seq resolves transcriptome signatures within the spatial context of intact tissue by integrating histology with RNA-seq [14,15]. Both histology and RNA-seq are completed in a sequential manner on the same tissue section placed on a micro-arrayed glass slide [14,16–18]. ST-seq begins with the histology component, involving fixation, H&E staining and imaging. The subsequent RNA-seq component begins with the release of RNA from the intact tissue section for capture by arrayed oligo-dT spots, termed ST-spots, which also contain a spatial barcode. Each ST-spot captures transcriptome information from one to nine adjacent cells, depending on the slide technology and the tissue type. The captured polyadenylated RNA is reverse transcribed to cDNA with the spatial barcode, then denatured and processed for library preparation and sequencing. The sequenced spatial barcode is then used to map the captured RNA to an ST-spot. Then the ST-spots are aligned with the H&E image to visualise the transcriptome-wide gene expression within the spatial context of the intact tissue. Currently ST-seq has been used in embryonic, inflammatory and cancer tissue, but has yet to be extended to the mammalian kidney [14,18–29].

In this study, we used a commercially available ST platform to investigate spatially resolved transcriptome expression in healthy human and mouse kidney tissue. We generated an ST profile of the mammalian kidney, allowing spatially resolved differential gene expression (DGE) analysis, spatial identification of functional nephron segments, cell-to-cell interaction (CCI) analysis in glomeruli, and chronic kidney disease associated genetic variant calling. All newly-generated ST-seq data from the human and mice kidneys has been deposited in a public repository (address).

## Results and Discussion

Frozen 10 μm sections from four human cortical kidney tissues (Patients A-D) and six whole mouse kidneys were processed for ST-seq. **Fig. 1.a** demonstrates the generation and analytical workflow of the ST-seq data from the mammalian kidneys.

**Fig. 1:**
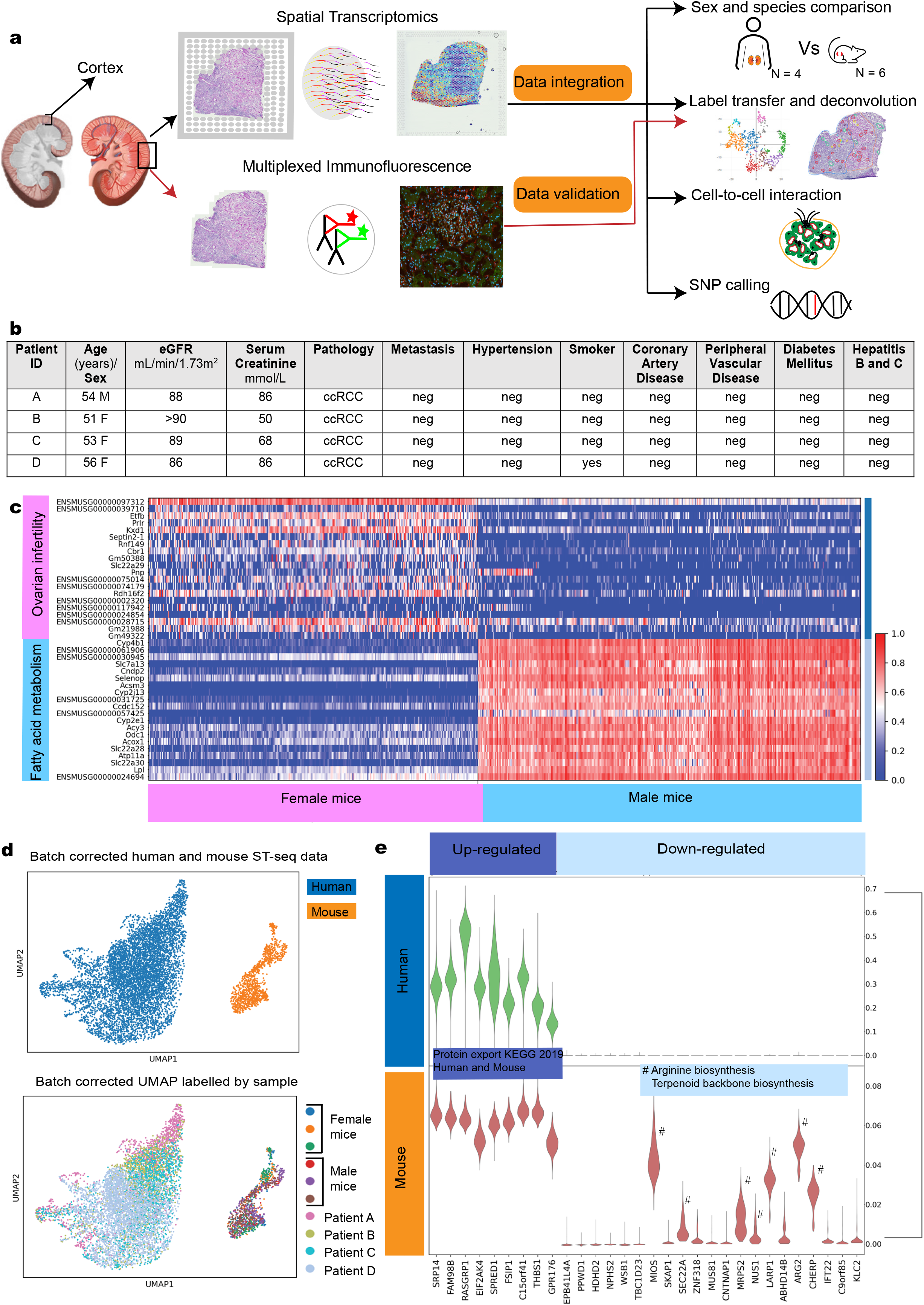
Spatially resolved transcriptome profiling in mammalian kidney. **a** A schematic of the workflow for generation and analysis of ST-seq data from mammalian kidneys. **b** Patient cohort characteristics table. **c** Heat map illustrating the top 40 DE genes, between the sexes in mice cortical kidney tissue regions. **d** Integration of human and mouse cortical kidney tissue data; (top panel) UMAP showing the batch corrected human and mouse ST-seq data clustered separately for the two species, (bottom panel) Sample labelling of the batch corrected UMAP shows an integrated heterogeneous human and mouse kidney samples, in their respective species cluster showing no batch effect. **e** Violin plot illustrating the top 30 DE genes, identified by DGE, between the human and mouse cortical kidney tissues.

Human kidneys were from one consenting male (Patient A) and three female (Patients B to D) patients that were matched for comorbidities and aged 51 to 56 years old **(Fig. 1.b).** ST-seq of the cortical region of the human kidney collectively detected over 23,000 genes (GRCh38-3.0.0) **(Fig. S1.a)**. Mouse kidneys were collected from 6-8 week old (C57BL/6J, wild type) mice. Collectively the ST-seq within the mouse kidney detected over 22,000 genes (GRCm38 - mm10) **(Fig. S1.b)**.

In the human ST-seq data, we identified high levels of mitochondrial RNA (mtRNA) expression **(Fig. S2)**. This observation is most likely due to the high metabolic requirements of mammalian kidneys to perform primary homeostatic functions, compounded by an enriched capture of mtRNA due to their polyadenylation [30–34]. The top 10 most highly expressed genes in all four human ST-seq data, were enriched for ATP synthesis coupled electron transport (GO:0042775) and respiratory electron transport chain (GO:0022904) networks, consistent with high energy requirements. Therefore, for human ST-seq data, we used a high threshold to filter only those ST-spots where mtRNA represented at least 50% of total reads **(Fig. S2)**.

Next, we performed DGE analysis of human and mouse (cortical) ST-seq data to explore quantitative changes in spatially resolved transcriptomes between sexes and species. We first completed batch correction and integration of all mouse ST-seq data to remove non-biological variability from our ST-seq data [35] **(Fig. S3.a-d)**. We performed DGE analysis between sexes with the mouse ST-seq data, demonstrating that the top 40 differentially expressed (DE) genes between sexes separated the female and male mice. Further enrichment analysis of these DE transcripts identified genes associated with fatty acid metabolism in male mice and genes associated with ovarian infertility in the female mice **(Fig. 1c)**. We subsequently performed DGE analysis in the human ST-seq data between sexes, but found no DE genes. We attributed this to the fact we were limited to one male sample, which restricted DGE analysis between sexes in the human ST-seq data.

We next conducted DGE analysis between species in our ST-seq data. Gene expression studies of human and mouse kidneys have been extensively performed via bulk RNA sequencing [13,36,37]. However, the data do not extend to understanding the differences in spatial patterns between the kidneys of these two species. We first identified human orthologues of the mouse genes and used these orthologues for downstream analysis. We completed batch correction and integration with the human and mice (cortical) ST-seq data. We found no technical variation between the sexes in either mice and humans, however there was a marked separation between the species in the UMAP plot **(Fig. 1d)**. We found 30 significantly DE genes between species, including an enrichment for the protein export pathway with nine genes more highly expressed in human kidney samples (KEGG 2019; *SRP14, FAM98B, RASGRP1, EIF2AK4, SPRED1, FSIP1, C15orf41, THBS1, GPR176*) **(Fig. 1e)**. We also found seven genes involved in amino acid metabolism with higher expression in the mouse data *(MIOS, SEC22A, MRPS2, NUS1, LARP1,ARG2, CHERP*). However, the overall low number of detected DE genes indicates that mammalian kidneys, including both human and mouse cortical tissue, have highly similar transcriptome profiles.

From here we focused on the human ST-seq data to investigate cell types, their transcriptional signatures, and spatial locations, using two complementary analytical strategies - Seurat [38] and stLearn [39]. We initially defined the spatial organisation of the human kidney using Seurat and stLearn clustering to identify ST-spots with distinct transcriptome profiles and mapped these cluster identities to the H&E tissue images. We then tested this approach by identifying ST-spot clusters enriched for glomerular and vascular markers, mapping these to the H&E image. The presence of glomerular and vascular structures at the corresponding tissue location was validated and annotated by an expert pathologist, then correlated with multiplexed immunofluorescence (mIF) for protein markers of these cell types on consecutive tissue sections **(Fig. 2a)**. Clusters annotated in the H&E and mIF images correlated with both Seurat and stLearn clusters, specifically for glomeruli and large blood vessels. We performed Wilcoxon statistical tests to confirm the identity of the putative glomerular clusters, identifying established marker genes for the glomerulus within the top 20 DE genes, including NPHS2, PODXL and PLA2R [40]. Further gene enrichment analysis of the DE genes within the glomerular cluster identified functionally-relevant structures like ‘slit diaphragm’, a specialised intercellular junction between the foot processes of epithelial cell (termed podocytes) in the glomerulus [41,42] **(clusters marked as red in Fig. 2a)**.

**Fig.2:**
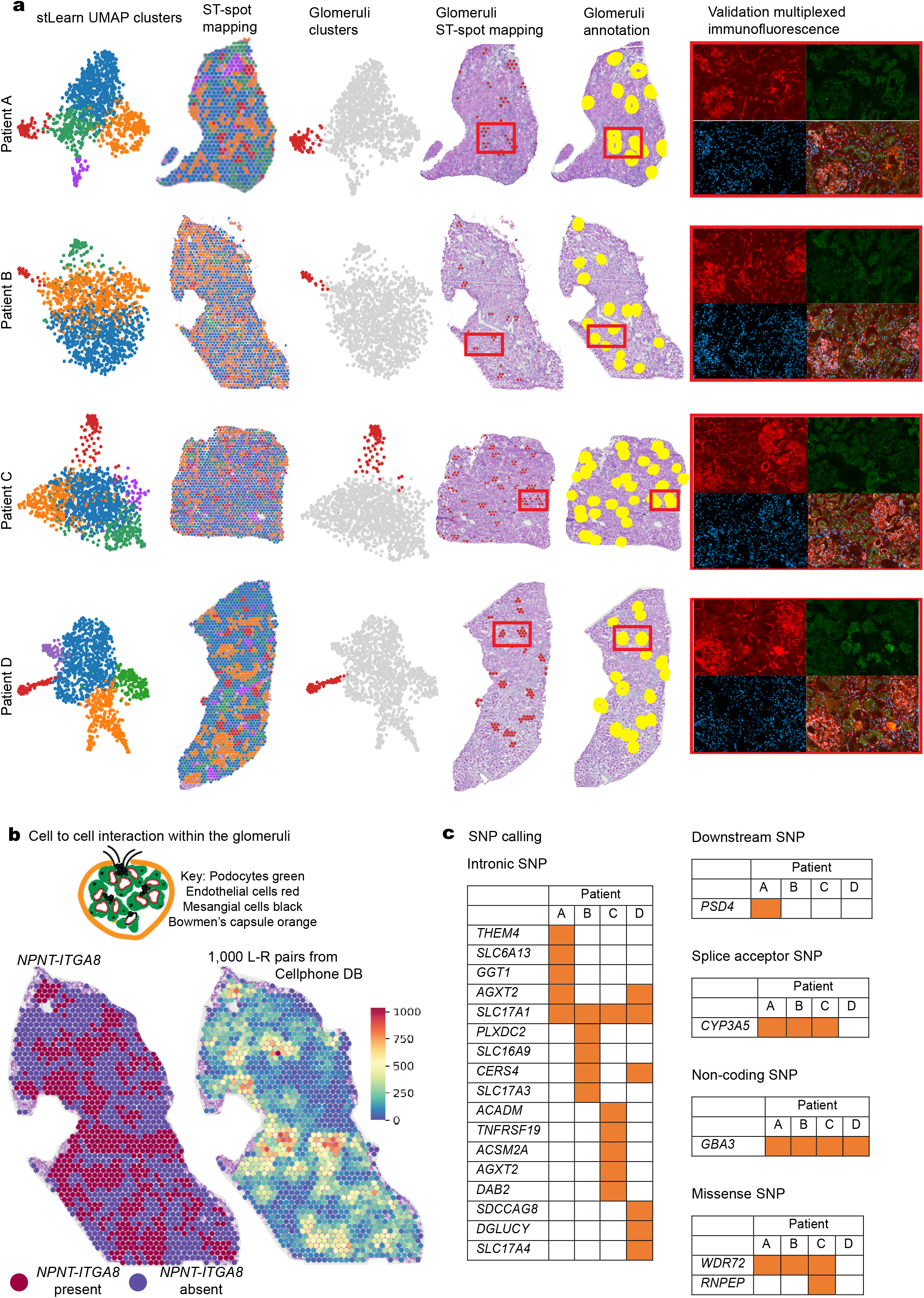
Integrative analysis of morphology, spatial expression, cellular interactions and genotypic effects in the generated human kidney ST-seq data. **a** Analysis of the human kidney ST-seq data with stLearn. From left to right; stLearn UMAP clusters, clustered ST spots mapped to the H&E image, glomerular clusters (as red) within the UMAP, glomerular clusters mapped to the H&E image, glomeruli annotated in the H&E images, and multiplexed immunofluorescence (mIF) staining; (top left) red demonstrates anti-CD31 staining for endothelial cells in the glomeruli and blood vessels, (top right) green demonstrates anti-AQP1 staining for identifying proximal tubular cells within the tubular compartment, (bottom left) blue demonstrates DAPI nuclear staining and (bottom right) multiplexed channel demonstrates glomeruli, vessels and proximal tubules. **b** Cellular communication via cell to cell interactions in the glomeruli are demonstrated in Patient B; (top panel) A schematic of a glomerulus, (bottom left) spatial expression of NPNT-ITGA8 (red is presence) in the glomeruli are mapped to the H&E image and (bottom right) the tissue interaction landscape of 1000 ligand-receptor(L-R) pairs (colour gradient shows interaction scores from randomness as 0 to significance as larger than 2). **c** A table of the 21 single nucleotide polymorphisms (SNPs) associated with chronic kidney disease identified in our human kidney ST-seq data.

In the human ST-seq data, we assigned ST-spots to key functional structures by applying Seurat’s label transfer functionality to identify proximal tubules, distal convoluted tubules, collecting ducts, Loop of Henle, interstitium and immune cells **(Fig. S2, two right-most columns)**. However, we noted that structures within the kidney, such as glomeruli, are composed of multiple different cell types, which may be captured within a single ST-spot. Additionally, we also had to account for the possibility that each human kidney ST-spot contained cell types from multiple functional structures. This concept was confirmed by assessment of the H&E images, which showed abutting discrete functional structures overlaying individual ST-spots. To account for this, we performed deconvolution, whereby the expression profile from thousands of genes detected in each ST-spot is compared to the expression profiles of cell type specific marker genes from a reference dataset, to predict the proportion of different cell types present in each ST-spot. Using SPOTlight [43], we performed deconvolution and identified the proportion of specific cell types (including dispersed immune cells) within each ST-spot **(Fig. S3c)**, providing higher resolution of the spatial localisation of kidney structures.

We extended our human kidney ST-seq data analysis to explore cellular communication between glomerular cells using a CCI algorithm [39]. Structurally, a glomerulus is a tuft of capillaries composed of mesangial, endothelial and podocyte cells, surrounded by the Bowman’s capsule lined with parietal epithelial cells [3,44]. Cell communication between podocytes and mesangial cells was explored as ligand-receptor (L-R) gene co-expression within ST-spots using stLearn **(Fig. 2b).** First we tested 20 published L-R pairs, including the nephronectin (*NPNT*)-integrin α8β1 *(ITGA8*) axis which governs mesangial cell behaviour [45]. Genes encoding both proteins were co-expressed in several ST-spots which included a broader region than identified by the glomerular clusters earlier. Next, we tested >1000 L-R pairs curated in the CellphoneDB database [46,47]. By applying stLearn [39], we mapped L-R gene spatial co-expression within and between ST-spots of glomeruli regions, identifying hundreds of putative interactions. We believe this CCI approach within the spatial context of functional structures and their cell types is crucial in understanding the molecular mechanisms of kidney physiology and pathology [48–50].

To explore the utility of the ST approach in human disease, we identified single nucleotide polymorphisms (SNPs) present in our human ST-seq data that are associated with chronic kidney disease in existing GWAS databases [51]. Across the four human kidney tissue sections, we identified >130,000 high confidence SNPs (p-value <0.01 or Phred quality scores >20). Among these SNPs, we found 36 SNPs overlapping reported CKD associated SNPs, 21 of which were high quality (probability of true SNP calling positive >0.99). These 21 SNPs were from intronic, missense, non-coding, downstream and splice acceptor sequences **(Fig. 2c)**. Of those 21 SNPs, 17 were from intronic regions of protein coding genes. Although RNA-seq strategies generally target processed RNA without introns, both scRNA-seq and ST-seq protocols utilise oligo-dT primers to capture polyadenylated sequences. These poly(A) tracts can lie within tails of mRNA, mtRNA, lncRNA or within intronic regions, resulting in some capture of intronic sequences [12]. We further visualised the spatial expression of the *SLC17A1* gene, which was associated with four detected SNPs in our ST-seq data (rs765285, rs1165151, rs1165213, and rs12212049) **(Fig. S3g)**. *SLC17A1* expression overlapped with proximal tubules in all our human kidney tissue sections, consistent with findings in previous studies [52].

## Conclusion

In this study, we have generated and analysed spatially resolved transcriptomes for human and mouse kidneys. The molecular expression profiles of these tissues were consistent with morphological annotations and molecular markers of key cell types, highlighting that ST-seq captures rich, anatomically meaningful biological information. This is demonstrated by our in-depth analysis of glomeruli, where stLearn clustering was used to identify ST-spots containing a glomerular gene expression signature. These ST-spots contained glomeruli as identified by histological and molecular markers. We demonstrated the utility of our analysis pipeline and the potential of these data resources to be used as a reference for a range of analyses, such as detection of GWAS-identified disease or trait associated genes and SNPs, comparison across sexes and species, and the analysis of complex CCI. This study lays a solid foundation for future studies using spatial transcriptomic data to investigate the mechanisms underlying mammalian kidney function under physiological and pathological conditions.

## Method

### Kidney tissue samples

In this study, we utilised healthy human cortical kidney tissues taken a minimum of 3 cm away from the tumour margins of four patients (3 females, 51 to 56 years old and 1 male, 54 years old). Tissue was collected for research purposes following informed patient consent and approval by the Royal Brisbane and Women’s Hospital Human Research Ethics Committee (2002/011 and 2006/072). During the collection of healthy human cortical kidney tissue, each patient was de-identified and their tissue snap frozen in standard biopsy cryomolds (Tissue-Tek, Sakura Finetek U.S.A) with optimal cutting temperature (OCT) compound (Tissue-Tek, Sakura Finetek U.S.A). We randomly allocated each patient a letter from A to D. This letter and corresponding non-identifying patient clinical information is provided in **Fig. 1b**.

The mouse kidneys utilised in this ST study were from three male (8 week old) and three female (6 week old) C57BL/6J wild type mice (Animal Ethics Committee approval UQDI/452/16 and IMB123/18). The mouse kidneys were collected during tissue harvesting and snap frozen in standard biopsy cryomolds (Tissue-Tek) with OCT compound (Tissue-Tek). These fresh frozen adult mouse kidneys were then stored at −80°C on site.

### RNA quality

Two 10 μm scrolls of tissue were collected in pre-chilled 1.5mL Eppendorf tubes from each frozen OCT block of healthy human cortical kidney (n = 4) and mouse kidney (n = 6) tissue. For each sample, RNA was extracted from the cryosectioned scrolls according to the QIAGEN RNeasy micro kit (Cat no: 74004), quantified according to the Qubit RNA HS assay kit (Cat no: Q32852) and the RNA integrity number (RIN) value assessed according to the Agilent 2100 Bioanalyzer RNA 6000 Pico assay (Cat no: 5067-1513). The measured RINs for all kidney tissues were greater than 7.

### Tissue optimisation

Tissue optimisation was performed according to the 10x Genomics ST Tissue Optimisation Manual (version 190219, 10x Genomics, USA) to determine the ideal permeabilization time. Frozen 10μm cryosectioned tissue from a healthy human cortical kidney and mouse kidney were utilised for this optimisation. The kidney tissue sections were dried at 37°C for 1 minute, fixed in pre-chilled 100% Methanol at −20°C for 30 minutes, stained in Mayer’s Haematoxylin (Dako) for 5minutes and Eosin (Sigma) for 2minutes. Imaging was performed on an Aperio XT brightfield slide scanner (Leica).

After H&E imaging the kidney tissue sections were placed in a permeabilization mix over a range of time points to allow the mRNA to drop down from the tissue sections and bind to the oligo-dT printed on the slide. The captured mRNA on the slide surface were then reversed transcribed to fluorescently labelled cDNA. This fluorescent cDNA signal was imaged on a Leica confocal microscope (SP8 STED 3X). The ideal permeabilization time was determined by correlating both the H&E and fluorescent images from the tissue optimisation slide. In this tissue optimisation slide the permeabilization time of 12 minutes generated the sharpest fluorescent signal that corresponded to morphological features noted in the H&E image. Hence a permeabilization time of 12 minutes was utilised for generating ST libraries for sequencing from human and mouse kidney tissue sections.

### Library preparation

ST library preparation of the healthy human cortical kidney tissues (n = 4) was performed according to the Visium Spatial Gene Expression Reagent Kits User Guide (CG000239 Rev C, 10x Genomics, USA). ST library preparation of the mouse kidney tissues (n = 6) was performed according to the ST Library Preparation Manual (version 190219, 10x Genomics, USA). In brief, 10μm cryosectioned human and mouse kidney tissues were placed onto respective pre-chilled library preparation slides. Sections were dried to the slides at 37°C for 1 minute, fixed in pre-chilled 100% Methanol at −20°C for 30 minutes, stained in Mayer’s Haematoxylin for 5 minutes and Eosin for 2 minutes. Brightfield imaging was performed on an Axio Z1 slide scanner (Zeiss). Based on the shorter (539 to 683bp) cDNA libraries generated from the healthy human cortical kidney tissue sections, we reduced the fragmentation reaction to 1 minute and the SPRI bead ratio was reduced to select for larger fragments. Then to further remove smaller library insert sizes (potentially consisting solely of TSO+poly(A)), we gel extracted the library preparations for patients A, B and C, followed by DNA clean-up according to Monarch PCR and DNA clean-up kit (Cat no: T1030S). All libraries were loaded at 1.8pM however, patients A, B and C, and mouse kidneys were sequenced using a High output reagent kit (Illumina), while patient D was sequenced using a Mid output reagent kit (Illumina), on a NextSeq500 (Illumina) instrument in-house at the Institute for Molecular Bioscience Sequencing Facility. Sequencing was performed using the following protocol: Read1 - 28bp, Index1 - 10bp, Index2 - 10bp, Read2 - 120bp.

### ST-seq library clean-up and mapping

Illumina generated ST-seq libraries, were first converted from raw base call (BCL) files to FASTQ files using bcl2fastq/2.17. Complex ST-seq libraries were retained and the FASTQ files were trimmed of poly-A sequences on the 3’ end and template switch oligo (TSO) sequences on the 5’ end using cutadapt/1.8.3 [53]. The cleaned FASTQ files were then mapped within Space Ranger V1.0 (10x Genomics) to the human reference genome and gene annotations (GRCh38-3.0.0) or mouse reference genome and gene annotation (GRCm38 - mm10). Finally, the genes were aligned to a resized H&E image from the library preparation based on the detection of the stained tissue and the fiducial markings.

### Multiplex immunofluorescence staining

Consecutive deeper 10 μm cryosections from the human cortical kidney tissues (n = 4) used for ST-seq, were placed onto room temperature SuperFrost® Ultra Plus slides (Thermo Scientific, U.S.A.). The tissue sections were then adhered to the slides by drying for 1 minute at 37°C and fixed with pre-chilled 100% methanol at −20°C for 30 minutes. Non-specific binding was blocked with 10% donkey serum (Merck-Millipore, Burlington,MA) for 15 minutes. Sections were incubated in a primary antibody mix comprising of anti-endothelial cell (monoclonal mouse anti-human CD31; Clone JC70A; Dako Omnis) and anti-Aquaporin-1 (polyclonal rabbit anti-human AQP1 (H-55); SC-20810; Santa Cruz Biotechnology) for 20 minutes. Fluorescent labelling was obtained with AlexaFluor conjugated secondary antibodies (donkey anti-mouse AlexaFluor PLUS 555 and donkey anti-rabbit AlexaFluor PLUS 488 (Invitrogen)) and DAPI (Sigma) mix incubated for 15 minutes. Slides were coverslipped with fluorescence mounting medium (Agilent Technologies, Santa Clara, CA). Imaging was performed on an Axio Z1 slide scanner (Zeiss) at 20x objective with Cyanine 3 (567nm), FITC (475nm) and DAPI (385nm) fluorescent channels. Image acquisition and analysis were performed within ZEN software (ZEN 2.6 lite; Carl Zeiss). Annotation of specific functional structures seen in the H&E image from the library preparation slide was correlated against the deeper consecutive multiplexed immunofluorescence image of the healthy human cortical kidney tissue sections.

### Seurat analytical pipeline

Human ST-seq data were demultiplexed using Loupe Browser (v4.0, 10x Genomics, USA) and were analysed using a modified version of the Seurat Spatial workflow (https://satijalab.org/seurat/v3.2/spatial_vignette.html). Preliminary quality control steps involved the filtration of spots containing more than 50% mitochondrial genes or 50% ribosomal genes; however, no samples reached this ribosomal threshold. 2000 variable features were detected by Seurat, and data were normalised using Scran [54] prior to running PCA analysis in Seurat. UMAP dimensionality reduction and clustering were performed using the first 50 principal components. Clustering was tested using a range of resolution values from 0.1 to 1.6 and the highest average stable resolution value was selected for each sample using the SC3 measure from Clustree [55]. The generated clustering results were visualized in both two dimensional UMAP space and in spatial context mapped over the H&E images.

We performed label transfer in two sequential steps using publicly available human kidney snRNA-seq [6] and scRNA-seq [12] datasets. This label transfer method projects known reference datasets and unknown datasets into a shared low-dimensional space, where equivalent cell types are arranged in the same neighbourhood in the two dimensional UMAP space, allowing for inference of cell types in the query dataset from the reference dataset. First, label transfer annotation from the snRNA-seq dataset was used to determine high-confidence ST-spot annotations. In the second round, the scRNA-seq data was used to label the remaining unlabelled ST-spots. In both rounds, transfer of cell type annotations from the reference to a query ST-spot was made if the confidence score for the top match was greater than 0.6; remaining ST-spots were left unannotated.

### stLearn analytical pipeline

The generated human and mouse ST-seq data was also analysed using stLearn, a novel Python-based toolkit [39]. stLearn uses the morphological similarity between neighbouring ST-spots to normalise gene expression and reduce “dropout” noise, an inherent technical limitation of RNA-seq technologies [39,56,57]. With the mouse kidney ST-seq data, we first filtered out genes that were expressed in less than 3 ST-spots. The filtered gene count matrix was then normalised by counts per million method, followed by log transformation and scaling. Finally, tissue morphological information was used to normalise the gene count matrix by running the stSME function. Downstream clustering analysis identified three clusters in the female mice that defined the spatially functional regions of the cortex and both outer and inner medulla. In the male mice, stLearn identified two clusters that defined the spatially functional regions of the cortex and medulla. The presence of the inner medulla cluster was attributed to the depth of the tissue sections which was greater in the female than the male mice kidneys. Hence for DGE analysis, between sexes and species, we selected the cortical region in both male and female mice.

With the human kidney ST-seq data, we first filtered the mitochondrial genes based on the quality of the data. In higher quality ST-seq data, we retained mitochondrial genes, whilst in lower quality ST-seq data, we removed all the mitochondrial genes. Additionally, we filtered ST-spots containing more than 50% mitochondrial genes. ST-spots with high total read counts relative to the total number of detected genes per spot were also filtered. The top genes based on expression levels were selected by using Scanpy [58], and the data were scaled to perform PCA analysis. Normalization (spatial smoothing step) which integrates gene expression profiles with tissue morphology using deep learning, was performed using the first 25 principal components. Leiden clustering was used to perform clustering analysis with flexible parameters. We used SPOTlight [43] to deconvolute the mixture of cell types in each spot. The same scRNA-seq [12] and snRNA-seq [6] datasets used for Seurat label transfer were also used for SPOTlight deconvolution.

### Analysis of GWAS single nucleotide polymorphisms (SNPs)

SNP variant calling was performed within the short sequence reads of ST-seq data generated from the healthy human cortical kidney tissues (n = 4) using FreeBayes [51], a Bayesian genetic variant detection program. From each of the mapped BAM files for human kidneys, we detected 140,241, 138,227, 150,935, and 205,471 high confidence SNPs respectively. We collected all reported SNPs associated with kidney diseases in the GWAS catalog database. SNPs called from ST-seq data were then compared to the known disease-associated SNPs. Expression values of those genes were visualised on the tissue sections.

## Supplementary figures

**Fig. S1:**
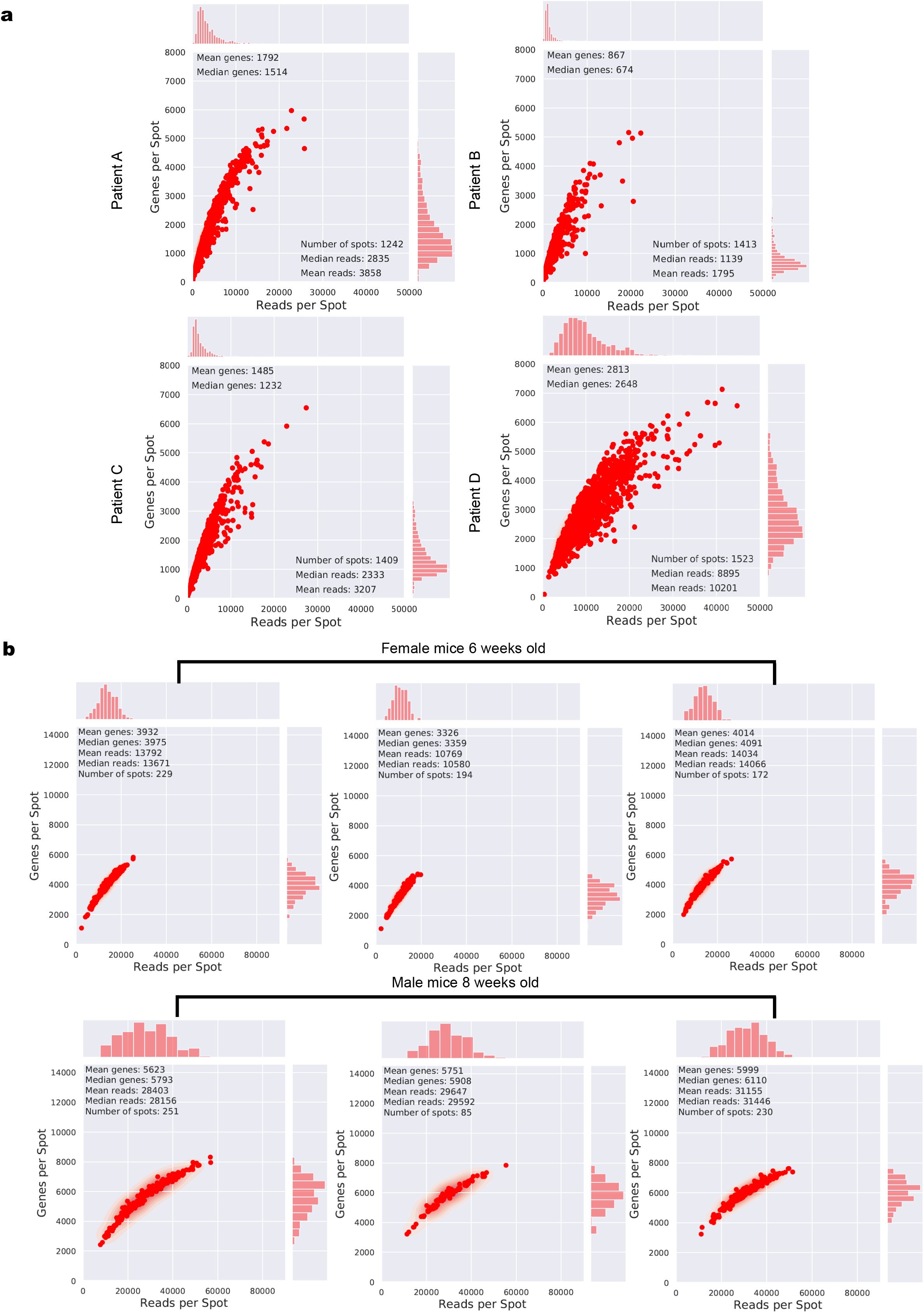
ST-seq data summary. **a** Scatter plots showing the number of genes (y-axis) against reads per spot (x-axis) detected in our ST-seq data for human kidneys. **b** Scatter plots showing the number of genes (y-axis) against reads per spot (x-axis) detected in our ST-seq data for mouse kidneys.

**Fig. S2:**
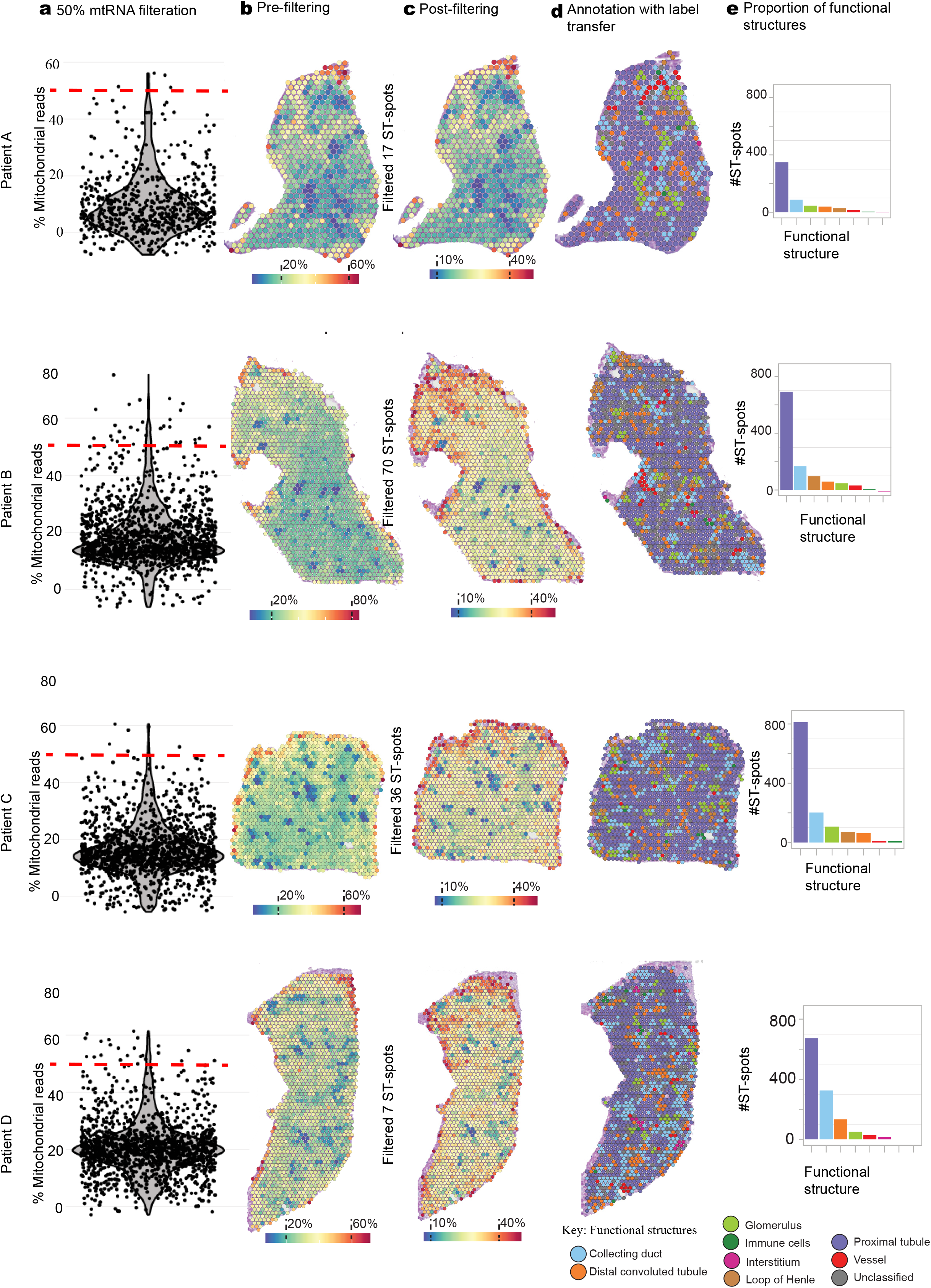
Analysis of human kidney ST-seq data with Seurat. **a** Violin plots showing the percentage of reads mapping to mitochondrial genes for each spot per sample. The red dashed line indicates the 50% threshold for filtering spots with high mitochondrial counts. **b-c** Spatial visualisation of mitochondrial read percentage values before (b) and after (c) filtering spots with >50% mitochondrial reads. **d** Annotation of spots by stepwise label transfer from snRNA-seq [6] and scRNA-seq [12] healthy human kidney reference datasets. Spots are coloured based on the highest-confidence functional annotation. **e** The number of spots assigned to each functional annotation. Bars are coloured by the annotation as in **d**.

**Fig. S3:**
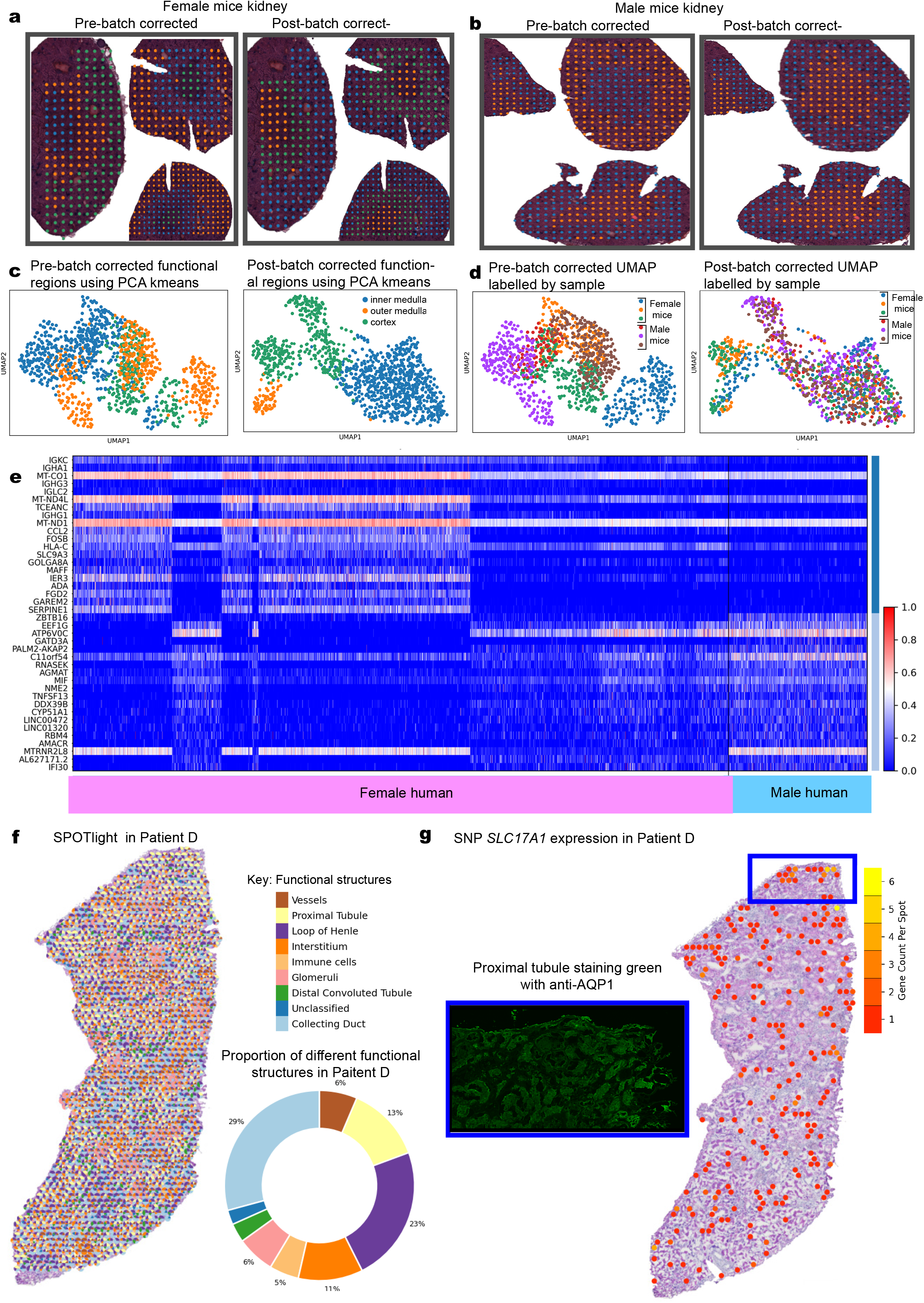
Analysis of the mouse kidney ST-seq with Seurat. **a** Mapping of the stLearn identified clusters to the female mice H&E images (left panel represents pre-batch and right panel represents post-batch corrected). **b** Mapping of the stLearn identified clusters to the male mice H&E images (left panel represents pre-batch and right panel represents post-batch corrected) **c** UMAP illustrating the pre-batch corrected (left) and post-batch corrected functional regions (right). **d** UMAP illustrating the pre-batch corrected (left) and post-batch corrected (right) functional regions labelled by each sample identity. **e** Heat map illustrating the top 40 differentially expressed genes between female and male human cortical kidney tissue sections. **f** SPOTlight analysis illustrating the deconvolution of key functional structures in Patient D. The pie chart demonstrates the proportion of these key functional structures in the cortical kidney tissue section of Patient D. **g.** Visualisation of gene expression for *SLC17A1,* identified as a target gene with the highest number of disease-associated SNPs detected in the ST-seq dataset across all four patients. These chronic kidney disease associated genetic variants include rs765285, rs1165151, rs1165213, and rs12212049, respectively. Multiplexed IF image for APQ1, reveals the region of the tissue expressing *SLC17A1* gene variants is predominantly proximal tubules.

## Declarations

### Ethics approval and consent to participate

The study used cortical kidney tissue samples from macroscopically healthy regions of renal cell carcinoma nephrectomies, following approval from the Royal Brisbane and Women’s Hospital Human Research Ethics Committee (Reference Number 2002/011; Approved 4 November 2016).

The study used whole mouse kidneys from C57BL/6J wild type mice, following approval from the University of Queensland Animal Ethics Committee (UQDI/452/16 and IMB123/18).

### Consent for publication

Informed consent from all patients was obtained during the pre-admission clinic in accordance with the Declaration of Helsinki as outlined in the approval from the Royal Brisbane and Women’s Hospital Human Research Ethics Committee (Reference Number 2002/011; Approved 4 November 2016).

### Availability of data and materials

The human and mouse kidney ST-seq datasets and codes are publicly available here at *https://github.com/BiomedicalMachineLearning/SpatialKidney.*

### Authors’ contributions

A.M.R., A.C., A.J.K., H.H., Q.N, and A.J.M conceived and designed the study; A.M.R., P.N.L, S.Y., S.M.T., S.E.H., J.C., and S.A. carried out the experiments; A.M.R., M.S,Y.N., A.J.K., H.H., and A.J.M. reviewed the patient data; D.P., X.T., L.F.G., and Q.N. performed the bioinformatics analyses; A.M.R., A.J.K., A.S., and L.F. performed the histological examination of the kidney; A.M.R., D.P., X.T., L.F.G., A.J.K., H.H., Q.N., and A.J.M. drafted the paper. All authors revised and approved the final version of the manuscript.

## Acknowledgements

The authors would like to thank the tissue donors, Queensland Health clinicians, pathologists and scientists, Conjoint Internal Medicine Laboratory for their support and discussions. The authors would like to thank the Australian Cancer Research Foundation (ACRF)/Institute for Molecular Bioscience (IMB) Cancer Biology Imaging Facility (established with the support of the ACRF), the UQ School of Biomedical Sciences Imaging Facility and UQ IMB Sequencing Facility for helpful discussions and guidance. The authors would like to thank Ronan Kapetanovic (IMB) and Ian Frazer (University of Queensland Diamantina Institute) for providing the mouse kidney tissues.

